# Fast and accurate spike sorting in *vitro* and *in vivo* for up to thousands of electrodes

**DOI:** 10.1101/067843

**Authors:** Pierre Yger, Giulia L.B. Spampinato, Elric Esposito, Baptiste Lefebvre, Stéphane Deny, Christophe Gardella, Marcel Stimberg, Florian Jetter, Guenther Zeck, Serge Picaud, Jens Duebel, Olivier Marre

## Abstract

Understanding how assemblies of neurons encode information requires recording large populations of cells in the brain. In recent years, multi-electrode arrays and large silicon probes have been developed to record simultaneously from hundreds or thousands of electrodes packed with a high density. However, these new devices challenge the classical way to do spike sorting. Here we developed a new method to solve these issues, based on a highly automated algorithm to extract spikes from extracellular data, and show that this algorithm reached near optimal performance both in *vitro* and *in vivo.* The algorithm is composed of two main steps: 1) a “template-finding” phase to extract the cell templates, i.e. the pattern of activity evoked over many electrodes when one neuron fires an action potential; 2) a “template-matching” phase where the templates were matched to the raw data to find the location of the spikes. The manual intervention by the user was reduced to the minimal, and the time spent on manual curation did not scale with the number of electrodes. We tested our algorithm with large-scale data from in *vitro* and in vivo recordings, from 32 to 4225 electrodes. We performed simultaneous extracellular and patch recordings to obtain “ground truth” data, i.e. cases where the solution to the sorting problem is at least partially known. The performance of our algorithm was always close to the best expected performance. We thus provide a general solution to sort spikes from large-scale extracellular recordings.

## Introduction

Throughout the brain, local circuits represent information using large populations of neurons [Buzsaki, 2010], and technologies to record hundreds or thousands of neurons simultaneously in the brain are therefore essential. One of the most powerful and widespread techniques for neuronal population recording is extracellular electrophysiology. Recently, newly developed microelectrode arrays have allowed recording the local voltage from hundreds to thousands of extracellular sites separated only by tenth of microns [Berdondini et al., 2005, Fiscella et al., 2012, Lambacher et al., 2004], giving an indirect access to large neural ensembles with a high spatial resolution. In these recordings, the spikes from each recorded neuron produce extracellular waveforms with a characteristic spatio-temporal profile across the recording sites. To access the spiking activity of individual neurons, we need to reconstruct the waveform produced by each neuron and tell when it appears in the recording. This process, called spike sorting, has received a lot of attention for recordings with a small number of electrodes. However, for large-scale and dense recordings, the problem of reliably extracting the spike contributions from extracellular recordings is still largely unresolved.

Current spike sorting algorithms cannot process this new type of data for several reasons. First, many algorithms do not take into account that the spikes of a single neuron will evoke a voltage deflection on many electrodes. Then, most algorithms do not scale up to hundreds or thousands of electrodes in *vitro* and *in vivo,* because their computation time would increase exponentially with the number of electrodes [Rossant et al., 2016]. Finally, all the solutions proposed so far usually require a significant amount of manual curation. Datasets from thousands of electrodes necessitate a spike sorting algorithm that is extensively automated. Furthermore, the few algorithms that have been designed to process large-scale recordings have not been tested on data where one neuron is recorded by the large-scale recordings and simultaneously by another technique, so that the success rate of the spike sorting algorithm can be measured [Pachitariu et al., 2016, Leibig et al., 2016, Hilgen et al., 2016].

Here we present a novel method for spike sorting that can scale up to recordings from thousands of electrodes. Based on a combination of density-based clustering and template matching, the method is fully automated, and only includes a final step of manual curation whose duration does not scale with the number of recorded cells. We tested the performance of the algorithm in two ways. First, we created several synthetic or “hybrid” datasets to properly quantify the performance of the algorithm. Then, we performed experiments where a large-scale extracellular recording was performed while one of the neurons was recorded with a patch electrode, and show that the performance of our algorithm was always close to an optimal classifier, both *in vitro* and *in vivo*. Therefore, our method appears to be a general, fast and scalable solution for spike sorting.

## Results

We sorted several datasets taken from large-scale in *vitro* and in vivo recordings. In all cases, electrodes were densely packed such that a spike from a single cell would affect the voltage of several electrodes. In vitro, we analyzed recordings from retinal ganglion cells recorded with planar multi-electrode arrays: rodent retina were recorded either by 252 electrodes spaced by either 30 or 60 microns, or by 4255 electrodes spaced by 16 microns. In vivo, we analyzed recordings of the cortex taken with 32 and 128 electrodes spaced by 20 microns (see Methods).

### Identifying templates for each cell

We developed an algorithm with two main steps, a clustering followed by a template matching step (see Methods for details). First, we detected the spikes as threshold crossings (fig. 1A), and isolated the extracellular waveforms for a number of randomly chosen spike times, that we call snippets in the following. Snippets were then clustered in different groups that should correspond to putative cells. Compared to classical sorting methods, here our purpose was not to obtain a full clustering of all the snippets, but rather to get the centroid of each cluster, which is way less demanding. We also made several natural assumptions so that this clustering step could be divided in parallel tasks that could be run independently.

**Figure 1:**
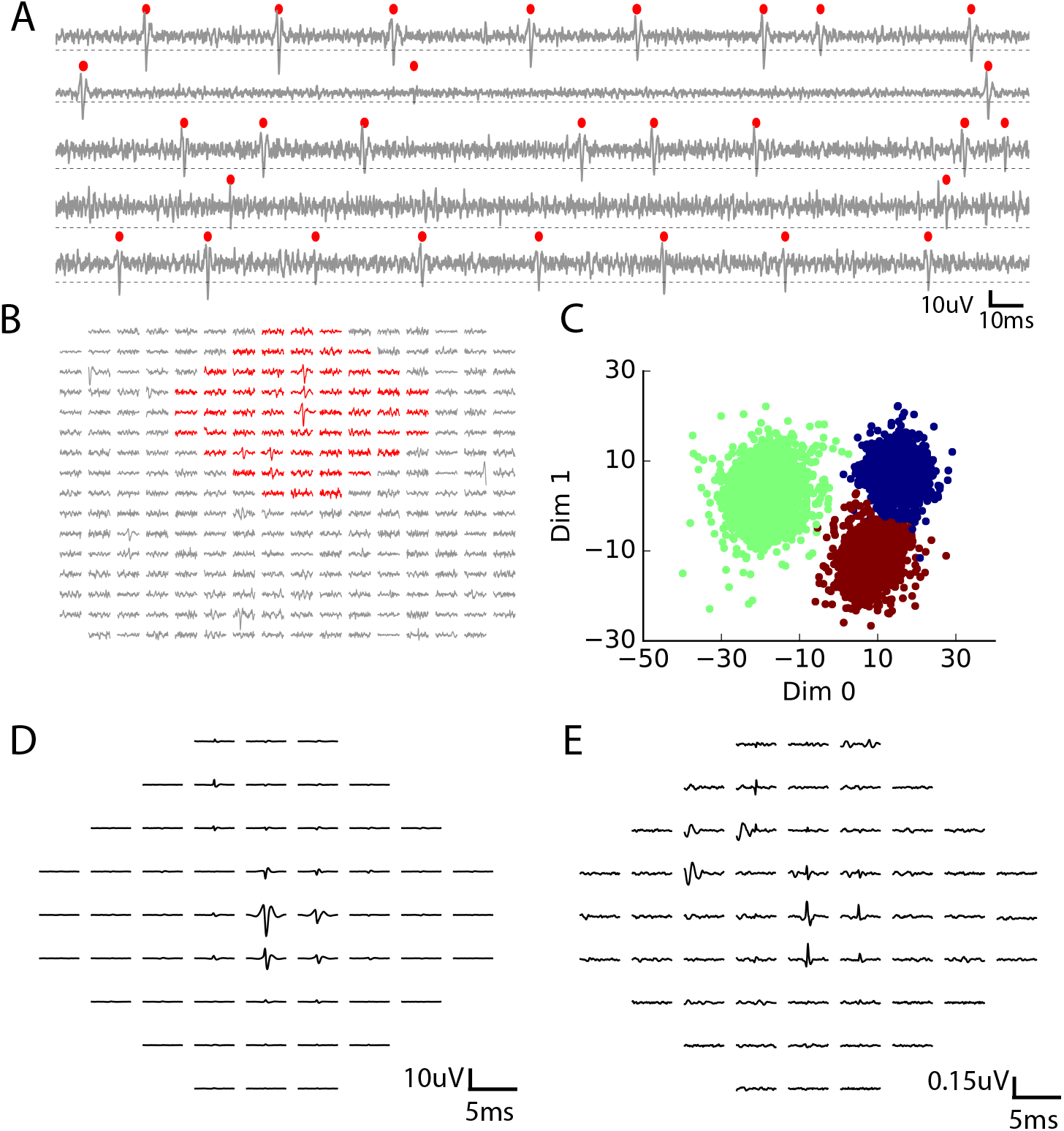
Local clustering of the snippets **A**. Five randomly chosen channels, each of them with its own detection threshold (dash dotted line). Detected spikes, as threshold crossings, are indicated with a red marker **B**. Example of a spike in the raw data. Red: electrodes that can be affected by the spike, i.e. the ones close enough to the electrode where the voltage peak is the highest; gray: other electrodes that should not be affected. **C**. For snippets collected in a given electrode, a robust density-based clustering detected three non-overlapping color-coded clusters (see Methods) **D**. The first component of the template ***T***(*t*), corresponding to the green cluster shown in **C. E.** The second component of the template ***U***(*t*) associated to the same green cluster (see Methods).

We divided the snippets in subsets according to their physical positions: we grouped together the snippets peaking on the same electrode. Following this we had as many groups of snippets as electrodes. Each group was then clustered independently. To efficiently cluster the snippets inside each of these groups, we had to reduce the dimensionality of the snippets. We assumed that a single cell can only influence the electrodes in its vicinity, i.e. close enough to its physical location (fig. 1B). We discarded all the electrodes whose distance to the peaking electrode was above a pre-defined radius (see fig. 1B). We also projected each snippet on stereotyped temporal waveforms to reduce the dimensionality (see Methods).

This subdivision in groups allows clustering each group in parallel, such that the computation time of this step only scales linearly with the number of electrodes, and can be divided by the number of computing cores available. We then performed a density-based clustering inspired by [Rodriguez and Laio, 2014] (see Methods) on each group. This density-based clustering is very efficient at finding the centroid of each cluster, which meets exactly our needs (see fig. 1C for three identified clusters on a given electrode). We adapted the algorithm to increase its robustness, and to avoid fine tuning parameters for a given dataset (see Methods). The resulting clusters were only used to define a “template” for each cell.

This template is a simplified description of the cluster and is composed of two waveforms. The first one is the average extracellular waveform inside the cluster (fig. 1D). The second is the direction of largest variance that is orthogonal to this average waveform (fig 1E; see Methods). We assumed that each snippet (i.e. voltage deflection) triggered by this cell is a linear combination of these two components. Finding these components for each cell does not require to have all the spikes of these cells clustered together: only a subset is good enough to estimate them. Therefore, this template extraction step works even if the clustering step only assigned a subset of spikes, which allowed processing very large datasets.

At the end of this first step, we have extracted an ensemble of templates (i.e. pairs of waveforms) that correspond to putative cells. By focusing on only getting the cluster centroids, we have made the clustering task easier. However, we have not assigned all the spikes to a cell. Therefore, we used a template matching algorithm to find all the instances where each cell has emitted a spike.

### Matching templates to the raw data

We assumed that the templates of different cells spiking together sum linearly and used a greedy iterative approach inspired by the projection pursuit algorithm (fig. 2A, see Methods). Within a piece of raw data, we looked for the template whose first component had the highest similarity to the raw signal and matched its amplitude to the signal. If this amplitude falls between predetermined thresholds (fig. 2A, B, C), we matched and subtracted the two components to the raw signal. We then re-iterated this matching process until no spike could be matched anymore (fig. 2D) (see Methods). In most cases, the spike times of a given template showed a nice refractory period (fig. 2E).

This template matching algorithm could be run independently on different blocks of data, such that the computing time only scaled linearly with the data length. We also took advantage of GPU computing resources to accelerate the computations. Each step of the spike sorting algorithm was parallelized, so the runtime of the full algorithm decreased proportionally with the numbers of CPU cores available (fig. 3A). As a result, the sorting could process one hour of data recorded with 252 electrodes in one hour with 9 CPU cores (spread over 3 computers) (fig. 3A, B). It also scaled up linearly with the number of electrodes (fig. 3B), and with the number of templates (fig. 3C). It was therefore possible to run it on long recordings (> 30min) with more than 4000 electrodes.

**Figure 2:**
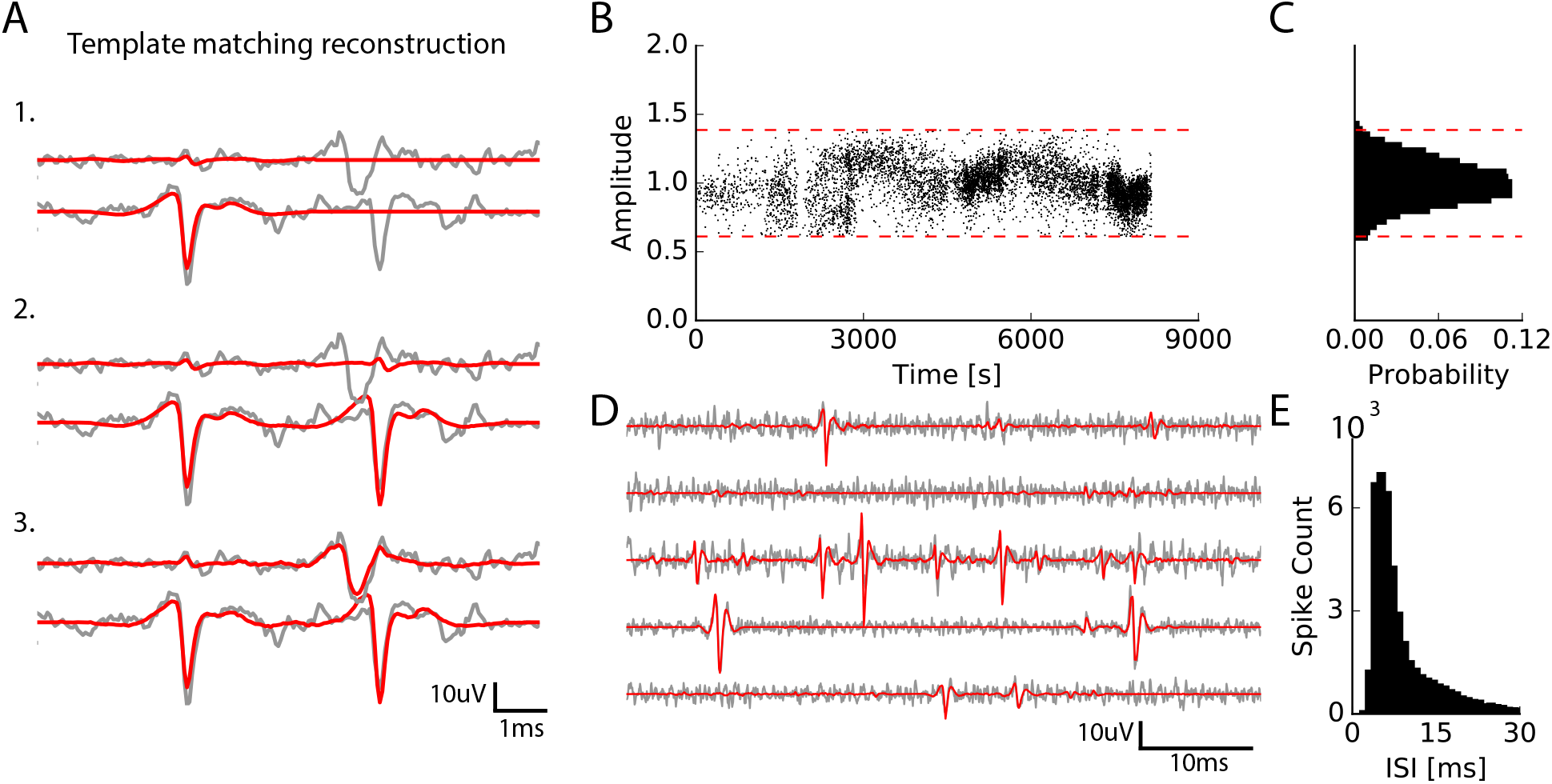
Illustration of the template matching phase **A**. Graphical illustration of the template matching with *in vitro* data (see Methods). Every line is a channel. Grey: the real data. Red: sum of the templates added by the template matching algorithm ; top to bottom: successive steps of the template matching algorithm (red). **B**. Examples of the fitted amplitudes for the first component of a given template, as a function of time. Each dot correspond to a spike time at which that particular template has been fitted to the data. Dashed dotted lines represent the boundaries allowed during the matching procedure. **C**. Distribution of the amplitudes for the same template, centered around 1. **D**. Final result of the template matching. Gray: extracellular signals for five channels. Red: sum of the matched templates. **E**. A typical Inter-Spike Interval (ISI) for a given template, showing a clear refractory period.

### Merging similar templates

After running our algorithm, despite a cleaning step where we removed obvious duplicates or sums of templates (see Methods), it is very likely that a single cell can be represented by two templates or more. Therefore, it is necessary to merge the templates and the corresponding spike trains that represent the same cell. Two criteria can be used to merge templates corresponding to the same cell. First, their templates should be similar. Second, there should be a dip around 0 in the cross-correlogram between the two spike trains (fig. 3D). This is due to the refractory period of the cell: two consecutive spikes from the same cell have to be separated by at least a couple of milliseconds.

To detect which pairs of spike trains had a dip in their correlogram, we computed the difference between the cross-correlogram of the two spike trains and a control, time-reversed correlogram over a given time window (fig. 3D; see Methods). This difference was plotted against the control value over the same time window for all the pairs with a high template similarity (fig. 3D). Almost all the pairs that needed to be merged appeared as an isolated group of points close to the equality line, which corresponds to a very strong dip in the cross-correlogram. To demonstrate this, we first simulated an ensemble of spike trains belonging to fake neurons and we randomly split them into several subsets (see Methods). This simulated data gave us a case where we know exactly which pairs need to be merged, and which ones correspond to different cells. We observed that all the pairs that had to be merged were grouped along the equality line, as expected (fig. 3D, left).

In a more realistic example, we took spike trains extracted from a large-scale recording where the merging was done manually (fig. 3E), and observed a similar grouping. In this representation, all the pairs that needed to be merged could therefore be selected all at once. We developed a graphical user interface to automatically plot these pairs, visualize individual cross-correlograms, and select the group of points of pairs that needed to be merged. The user could then quickly isolate and select the pairs of templates to be merged. By deciding which pairs needed to be merged from a single plot, this method avoids going through all the pairs to decide which ones need to be selected and drastically reduces the time of manual curation. This saves the time of the user, but also avoids having a criterion that is purely subjective and could change over the time spent at doing manual curation.

### Performance validation

We first validated our method with simulated ground truth datasets where artificial spikes were added in the recordings and retrieved by the algorithm. We ran the spike sorting on different datasets, picked some templates and used them to create new artificial templates, that we added at random places to the real recordings (see Methods and fig. 4A). We then run our sorting algorithm on this novel or “hybrid” datasets with these novel artificial cells randomly added, and measured if the algorithm was able to find at which times the artificial spikes were added. We counted a false negative error when such a spike was missed, and a false positive error when the algorithm detected a spike while there was not any (see Methods). Summing these two types of errors, the total error rate remained below 5% for all the spikes whose size was significantly above threshold. Fig. 4B shows the exhaustive results for 100 templates injected into a 252 electrodes recording in the retina, but the size or the type of the recording did not affect the error rates: they were similar for recordings with 4255 electrodes *in vitro* or 128 electrodes *in* vivo. Performance did not depend on the firing rate of the injected templates (fig. 4C), and only weakly on the normalized amplitude of the templates (fig. 4D), as long as it was above the spike threshold. Performance was not only satisfying for individual cells, but also to estimate correlations properly. We injected templates with a controlled amount of overlapping spikes (see Methods). The algorithm was always able to estimate the pairwise correlation between the spike trains (fig. 4E). The ability of our template matching algorithm to resolve overlapping spikes thus allowed an unbiased estimation of correlations between spike trains.

**Figure 3:**
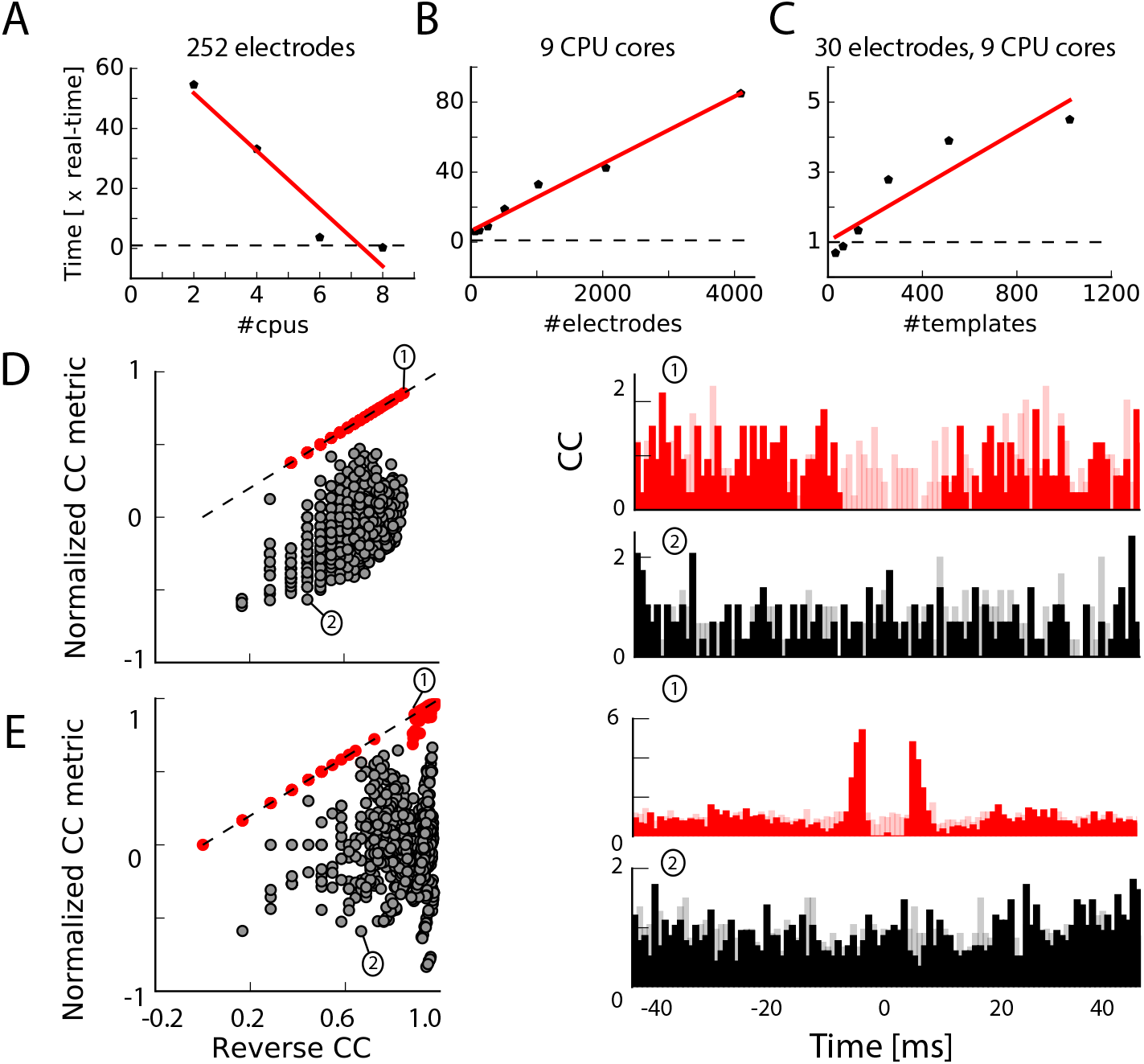
Management of large datasets **A**. Execution time as function of the number of processors, for a 90min dataset *in vitro* with 252 channels, expressed as a real-time ratio, i.e. the number of hours necessary to process one hour of data. **B**. Execution time as function of the number of channels, for a 30min dataset recorded *in vitro* with 4225 channels. **C**. Execution time as function of the number of templates, for a 10min synthetic dataset with 30 channels. **D**. Normalized Cross Correlation Metric compared to the Reverse Correlation for artificially generated and split spike trains (see Methods). Red: pairs of templates originating from the same neuron, that have to be merged. Black: pairs of templates corresponding to different cells. Insets on the right: for two chosen pairs (see numbers) the full cross-Correlogram (plain color) and the reverse correlogram (shaded color) **E**. Same as **D**., but with 10 real neurons selected after sorting a dataset recorded with 252 channels on a rat retina (see Methods).

**Figure 4:**
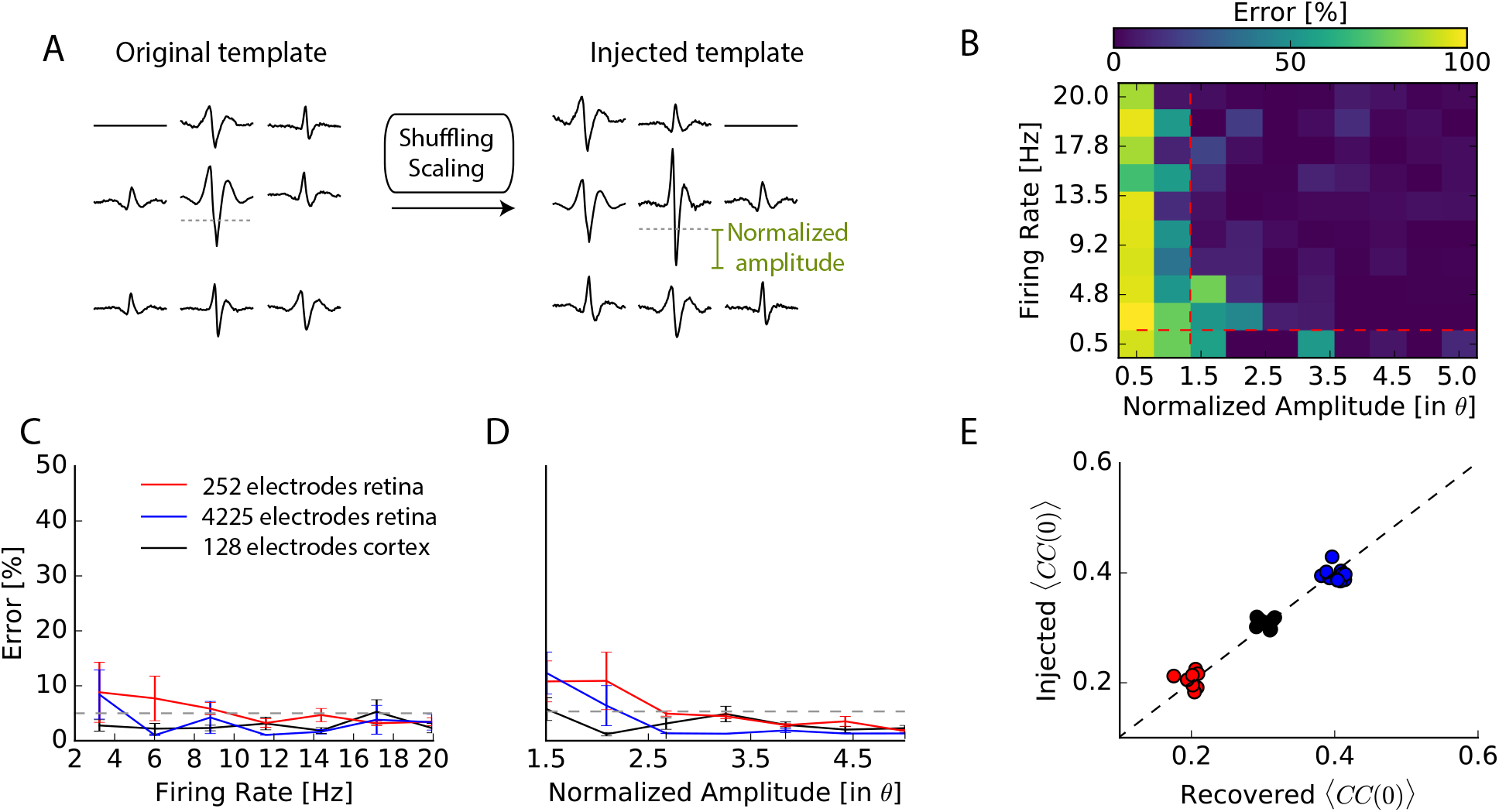
Performance of the algorithm on hybrid ground truth data **A**. A chosen template is injected elsewhere in the data as a new template. Dashed-dotted lines shows the detection threshold on the main electrode of the template, and normalized amplitude is expressed relatively to this threshold (see Methods). **B**. Error rates as function of the amplitude and the rate of injected templates, in a 252 electrode recording performed *in vitro* in the retina (see Methods). **C**. Mean error rate as function of the firing rate of injected templates, in various datasets. Errors bars show the standard error over 8 templates **D**. Error rate as function of the normalized amplitude of injected templates, in various datasets. Errors bars show standard error over 9 different templates E. Injected and recovered cross-correlation value between pairs of neurons for 5 templates injected at 10Hz, with a normalized amplitude of 2 (see Methods).

### Performance on ground truth data

The previous tests were done using “hybrid” data mixing real recordings and artificial templates. To test our algorithm in a more realistic situation, we performed dual recordings (fig. 5A, B). While we recorded many cells using a multi-electrode array (see schematic on fig. 5A), we simultaneously performed loose-patch recording of one of the cells, and recorded its spikes (fig. 5B). For one of the cells we know what should be the output of the spike sorting. In vitro, we recorded 26 neurons from 14 retinas with a 252 electrode MEA where the spacing between electrodes was 30 microns (see Methods). To test our algorithm in cases where the electrode density is lower, we generated datasets where we removed the signals of some electrodes, such that the density of the remaining electrodes was either 42 or 60 microns (by removing half or 3 quarters of the electrodes, respectively).

We then ran the spike sorting algorithm on the extracellular data, and estimated the error rates for the cell recorded in loose-patch, where we know where the spikes occurred. The performance of the algorithm will depend on the waveform triggered by the cell on the extracellular electrodes, as this is an intrinsic limit to the problem of spike sorting. If a spike of the patch-recorded cell triggers a large voltage deflection, this cell should be easy to detect. On the other hand, if the triggered voltage deflection was barely detectable, we expect that our algorithm should not perform well. To quantify this, we estimated a theoretical “best” performance for each recording. This best performance was found by training a non-linear classifier on the extracellular waveforms triggered by the spikes of the recorded cell, similar to [Harris et al., 2000, Rossant et al., 2016] (referred to as the Best Ellipsoidal Error Rate (BEER), see Methods). Note that this classifier “knows” where the true spikes are and simply quantifies how well they can be separated from the other spikes based on the extracellular recording. On the contrary, our spike sorting algorithm does not have access to any information about when the patch-recorded cell spiked.

We estimated the error made by the classifier and found that the performance of our algorithm almost always matched the performance of this classifier (fig. 5C, left), and this over a broad range of spike sizes. This suggests that our algorithm reached an almost optimal performance on this in *vitro* data. As expected, larger deflections lead to better performances (see inset of fig. 5C) for both the classifier and our algorithm, such that their respective performances were always close. Note also that because the BEER estimate is not designed to deal with overlapping spikes, it was possible for our template-matching based algorithm to obtain better performances in some cases.

We also used similar ground truth datasets recorded *in vivo* in rat cortex using dense silicon probes with either 32 or 128 recording sites [Neto et al., 2016]. With the same approach than for *in vitro* data, we also found that our algorithm reached a near optimal performance (fig. 5C, right). Together, these results show that our algorithm can reach an almost optimal performance (i.e. comparing to the BEER error) in various realistic cases, for both *in vivo* and *in vitro* recordings.

## Discussion

We have shown that a method based on density-based clustering and template matching allows to sort spikes from large-scale extracellular recordings both *in vitro* and *in vivo.* Our algorithm is entirely parallelized and could therefore easily handle long datasets recorded with thousands of electrodes. The only step that required manual intervention was the final step of merging spike trains corresponding to a same cell, that has been split in several templates by the algorithm. We presented a novel approach to merge all the spike trains at once during manual curation. This new procedure minimizes the time spent on manual curation. It also guarantees that all pairs will be merged according to similar criteria. We tested the performance of our algorithm on “ground truth” datasets, where one neuron is recorded both with extracellular recordings and with a patch electrode. We showed that our performance was close to an optimal nonlinear classifier that was trained using the true spike trains. Our algorithm has also been tested on purely synthetic datasets [Hagen et al., 2015], and similar results were obtained (data not shown).

**Figure 5:**
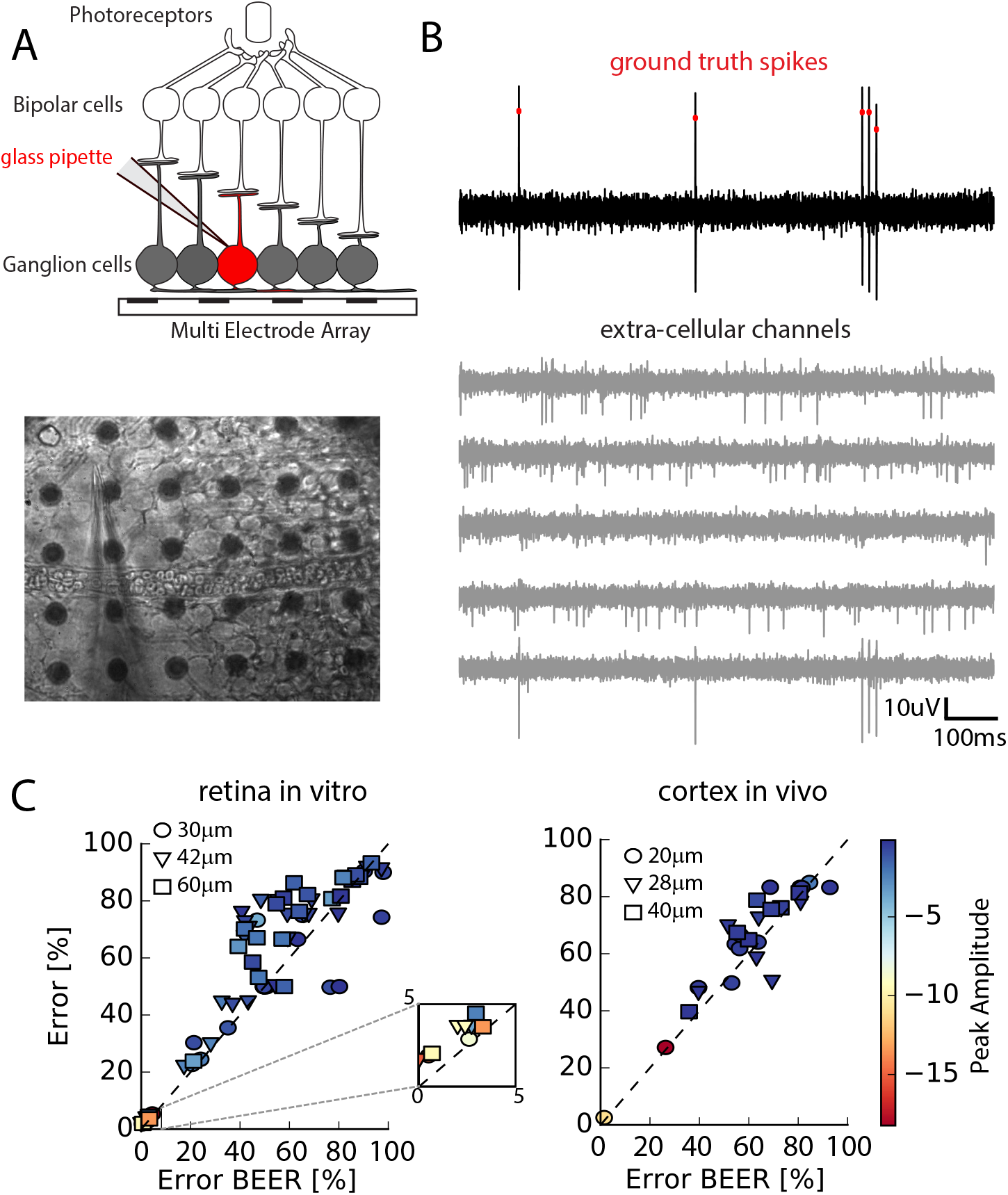
Performance of the algorithm on ground truth datasets **A**. Top: Schematic of the experimental protocol *in vitro.* A neuron close to the multi-electrode array (MEA) recording is recorded juxta-cellularly. Bottom: Image of the patch electrode on top of a 252 electrodes MEA, recording a ganglion cell. **B**. Top: Juxta-cellular recording showing the spikes of the recorded neuron. Bottom: Extra-cellular recordings next to the soma. **C**. Comparison between the error rates produced by the algorithm on different ground truth datasets and by the error rates of non-linear classifiers (Best Ellipsoidal Error Rate) trained to detect the spikes, either *in vitro* (Left), or *in vivo* (Right) (see Methods). Recorded neurons are color-coded as function of the maximal normalized amplitude recorded extra-cellularly.

Classical approaches to the spike sorting problem involve extracting some features from each detected spike [Hubel, 2015, Meister et al., 1994, Lewicki, 1994, Einevoll et al., 2012, Quiroga et al., 2004, Hill et al., 2011, Pouzat et al., 2002, Litke et al., 2004] and clustering the spikes in the feature space. In this approach, the spike sorting problem is reduced to a clustering problem, and this induces several major issues. First, to assign the spikes to the correct cell, the different cells must be separated in the feature space. Finding the exact borders of each cell in the feature space is a hard problem, that cannot be easily automated. As a result, most methods require a lot of manual curation, and therefore work only for a small number of electrodes. Second, running a clustering algorithm on data with thousands of electrodes is very challenging. Finally, overlapping spikes will appear as strong deviations in the feature space, and will therefore be missed in this approach. These three issues preclude the use of this approach for large-scale recordings with dense arrays of electrodes. In comparison, here we have parallelized the clustering step efficiently, and we have used a template matching approach, so that we only needed to get the centroid of each cluster, and not their precise borders. The template matching approach also allowed to deconvolve overlapping spikes in a fast and efficient manner.

Few template matching approaches have been tested, but only on in *vitro* data [Marre et al., 2012, Pillow et al., 2013]. Also, they only had one component for template matching, and did not allow any variation in the shape of the spike. Moreover, to get the templates, some used a manual approach [Segev et al., 2004, Prentice et al., 2011] that cannot be scaled up, or a k-means algorithm that would also be challenged by large-scale data. In all cases, they required a significant amount of manual curation, with a time that scaled linearly with the number of electrodes. Finally, they were not tested on in vivo data. in vivo recordings seem to be prone to more variation in the spike waveform than in *vitro* recordings, and we have designed our template matching method to take into account not only variation in the amplitude of the template, but also in shape. So our approach improves previous template matching algorithms by decomposing each spike as a sum of two waveforms.

Our method is almost fully automated. This paves the way towards online spike sorting for large-scale recordings. Several applications, like brain machine interfaces, or closed-loop experiments [Franke et al., 2014, Hamilton et al., 2015, Benda et al., 2007], will require an accurate online spike sorting. Our method is entirely parallel and can therefore be run in “real time” (i.e. one hour of recording processed in one hour) with enough computer power. This will require adapting our method to process data “on the fly”, processing new data blocks when they come, and probably adapting the shape of the template over time.

## Methods

### Experimental recordings

*in vitro* recordings with 252 or 4225 electrodes Retinal tissue was obtained from adult (8 weeks old) male Long-Evans rat (Rattus norvegicus) or mouse (mus musculus, 4-9 weeks old) and continuously perfused with Ames Solution (Sigma-Aldrich) and maintained at 32° C. Ganglion cell spikes were recorded extracellularly from a multi-electrode array with 252 electrodes spaced 30 or 60μm apart (Multi-Channel Systems), or with 4225 channels arranged in a 2D grid and spaced by 16μm [Zeck et al., 2011, Bertotti et al., 2014], at a sampling rate of 20kHz. Experiments were performed in accordance with institutional animal care standards.

For the ground truth recordings, electrophysiological recordings were obtained from *ex-vivo* isolated retinae of rd1 mice (4/5 weeks old). The retinal tissue was placed in AMES medium (Sigma-Aldrich, St Louis, MO; A1420) bubbled with 95% O2 and 5% CO2 at room temperature, on a MEA (10μm electrodes spaced by 30μm; Multichannel Systemps, Reutlingen, Germany) with ganglion cells layer facing the electrodes. Borosilicate glass (BF100-50, Sutter instruments) electrodes were filled with AMES with a final impedance of 6-9 MΩ. Cells were imaged with a customized inverted DIC microscope (Olympus BX 71) mounted with a high sensitivity CCD Camera (Hamamatsu ORCA −03G) and recorded with an Axon Multiclamp 700B patch clamp amplifier set in current zero mode.

For the data shown in fig. 2 and 4, we used a recording of 130min. For the data shown in fig. 5A, 16 neurones were recorded over 14 intact retinas. Recording durations ranged from 2min to 12min. The thresholds for the detection of juxta-cellular spikes were manually adjusted for all the recordings.

*in vivo* recordings with 128 electrodes We use the freely available datasets provided by [Neto et al., 2016]. Those are 32 or 128 dense silicon probes recordings (20μm spacing) at 30kHz performed in rat visual cortex, combined with juxta-cellular recordings. The dataset gave us a total of 13 neurons for fig 5. C, with recordings between 4 and 10min each. Similarly to the *in vitro* case, the detection thresholds for the juxta-cellular spikes were manually adjusted based on the data provided by [Neto et al., 2016] and on spike-triggered waveforms. For the validation with “hybrid” dataset, shown in fig. 4, we used the longest dataset recorded with 128 electrodes.

### Details of the algorithm

In the following, we consider that we have *N*_elec_ channels, acquired at a sampling rate *f*_rate_. Every channel *i* is located at a physical position ***p_i_*** = (*x_j_,y_j_*) in a 2D space (extension to 3D probes would be straightforward). The aims of our algorithm is to decompose the signal s over all channels 1,… *N*_elec_ as a linear sum of spatio-temporal kernels or “templates” (see equation 1).

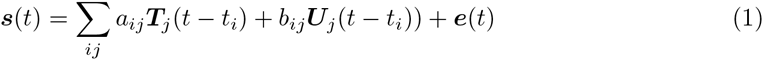

where *s*(*t*) is the signal recorded over *N*_elec_ electrodes and over multiple time points. ***T**_j_*(*t* – *t_i_*) and ***U**_j_*(*t* – *t_i_*) are the templates associated to each cell, which represents the waveform triggered on the electrodes by cell *j*. Times *t_i_* are all the putative spike times over all the electrodes. *a_ij_* and *b_ij_* are the amplitude factors for spike time *t_i_* for cluster *j*, and *e*(*t*) is the background noise.

The algorithm can be divided into two main steps, described below. After a preprocessing stage, we first run a clustering algorithm to extract a dictionary of “templates” from the recording. Second, we use these templates to decompose the signal with a template-matching algorithm. We assume that a spike will only influence the extracellular signal over a time window of size *N_t_* (typically 2ms for in vivo and 5ms for in *vitro* data), and only electrodes whose distance to the soma is below *r* (typically, 100μm for in vivo and 200μm for in *vitro* data). For every channel *i* centered on ***p_i_***, we define 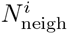 as the ensemble of nearby channels such that ||***P_i_*** – ***P_j_***||_2_ ≤ *r*. In the following, we note ***s**_i_*[*t*_min_, *t*_max_]_*i*_ the temporal slice of all time samples on channel *i* such that *t* ∈ [*t*_min_, *t*_max_].

### Pre-processing

#### Filtering

In a preprocessing stage, all the channels were individually high-pass filtered with a Butterworth filter of order three and a cutoff frequency of 500Hz, to remove any low-frequency components of the signals. We then substracted, for every channel *i*, the median such that ∀_*i*_, med(***s_i_***) = 0.

#### Spike detection

Once signals have been filtered, we computed a spike threshold *θ_i_* for every channel *s_i_* (*t*): *θ_i_* = *k*MAD(*s_i_*(*t*)), where MAD is the Median Absolute Deviation, and *k* is a free parameter. For all the datasets shown in this paper, we set *k* = 6. For every channel *s_i_* we detected the putative spike times 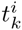 as all the local minima below *θ_i_*.

#### Whitening

To remove spurious spatial correlations between nearby recordings channels, we performed a spatial whitening on the data. To do so, we searched for a maximum of 20s of recordings where there are no spikes (i.e no threshold crossings). We then computed the Covariance Matrix of the noise ***C***_spatial_, and estimated its eigenvalues |*d_i_*} and associated eigenvectors {*V*}. From the diagonal matrix 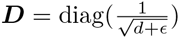, where *ϵ* = 10^−18^ is a regularization factor to ensure stability, we computed the whitening matrix ***W*** as ***VDV^T^***. In the following, each time blocks of data are loaded, they are multiplied by ***W***. After whitening, we recomputed the spike detection threshold *θ_i_* of each channel i in the whitened space.

#### Basis estimation (PCA)

To identify the spatio-temporal waveforms embedded in the data, we need to reduce their dimensionality. We collected up to *N_w_* spikes on each channel, at times 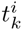. In order to compensate for sampling rate artifacts, we upsampled all the collected waveforms by bicubic spline interpolation to 5 times the sampling rate *f*_rate_, aligned on their local minima, and then re-sampled at *f*_rate_. We performed a Principal Component Analysis (PCA) on these centered and aligned waveforms and kept only the first *N*_PCA_ principal components. In all the calculations, we used default values of *N*_w_ = 10000 and *N*_PCA_ = 5. These Principal components were used during the clustering step.

### Clustering

The goal of the clustering step is to construct a dictionary of templates. As opposed to former clustering approaches of spike sorting [Quiroga et al., 2004, Harris et al., 2000, Kadir et al., 2014], because this clustering step is followed by a template matching, we do not need to perform the clustering on all the spikes: the templates are extracted from the centroids of the clusters.

#### Masking

We first collected many spikes to perform the clustering. To minimize redundancy between collected spikes, each time a local minimum 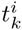 was selected on a given channel *i*, we prevented the algorithm to select any other minima in a spatio-temporal area around 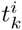, to avoid the selection of multiple threshold crossings originating from the same event. We excluded all the peaks on the neighboring channels 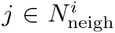 during a time window centered on 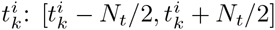.

#### Pooling of the spikes

In order to parallelize the problem, we used a divide and conquer approach [Marre et al., 2012, Swindale and Spacek, 2014]. Each time a spike was detected at time 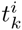 on electrode *i*, we searched for electrode *φ* where the voltage has the lowest value, i.e. such that 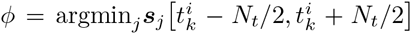. So for every electrode *i* we collected spikes peaking on this electrode. Each of these spikes is represented by a spatio-temporal waveform of size 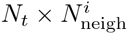. We projected each waveform on the PCA basis estimated earlier to reduce the dimensionality to 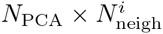. Note that during this projection, the same up-sampling technique described in the Pre-processing was used. For every channel *i* we collected 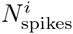 spikes, and each of them is a vector of size 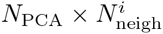. The maximal number of spikes collected is defined by the user as at *N*_spikes_, and we used a default value of *N*_spikes_ = 10000. To reduce dimensionality even further before the clustering stage, for every channel *i* we performed a PCA on the collected spikes, and kept only the first *N*_PCA_2__ principal components (in all the paper, *N*_PCA_2__ = 5). Therefore, we performed a clustering in parallel for every channel, on at max *N*_spikes_ described in a space of *N*_PCA2_-dimension. Note that this pre-grouping does not assume that the spikes are only detected on a single electrode. This clustering performed on each spike ensemble used the information available on all the neighboring electrodes.

#### Clustering by search of local density peaks

To perform the clustering, we used a modified version of the algorithm published in [Rodriguez and Laio, 2014]. For *N* data points ***x_i_***, *i* ∈ {1,…*N*} in a *M* dimensional space ℝ^*M*^ (in our case ***x_i_*** are spikes assigned to channel *i* and *M* = *N*_pca_2__), we computed the “density” *ρ* as the mean distance to the *S* nearest neighbors of ***x_i_***. *S* is chosen such that *S* = *ϵN*, with *ϵ* = 0.01. This density measure turned out to be more robust than the one given in the original paper, and rather insensitive to changes in *ϵ*. Then, for every point ***x_i_***, we computed *δ_i_* as the minimal distance to any other point ***x_j≠i_*** such that *ρ_j_* < *ρ_i_*. The intuition of the algorithm is that the centroids should be points with a high density (i.e. low *ρ*), and far apart from each others (high *δ*).

#### Accurate estimation of the density landscape

We performed the clustering only on a subset of 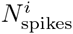 for each channel. To avoid potential inaccurate estimation of the densities *ρ_i_*, we collected iteratively additional spikes to refine this estimate. Keeping in memory the spikes *x_i_*, we searched again in the data 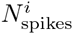 different spikes and used them to refine the estimation of *ρ_i_* of our selected points *x_i_*. When doing m passes, the complexity only scales as 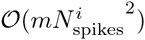. In all the following we used *m* = 3, and automatically stop updates if not enough spikes can be found in the dataset.

#### Centroids and cluster definition

To define the centroids we ranked the points as function of the products *ρδ*, we detected the best 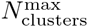 points as putative centroids. Clusters were formed by an iterative rule, going from the points of lowest rho to the points of highest *ρ*: each point was assigned to the same cluster than the closest point with a lower *ρ* [Rodriguez and Laio, 2014]. We created at max 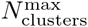 clusters. Once this is done, we iteratively merged pairs of clusters that were too similar to each others.

#### Merging similar clusters

We computed a normalized distance *ζ* between each pair of clusters *y*_1_ and *y*_2_. The center ***α_i_*** of cluster *Y_i_* was defined as ***α_i_*** = med(x_i_ ∈ **Y_i_**), and we can therefore project all the points from clusters *Y*_1_ and *Y*_2_ onto the axis joining the two centroids ***γ*_1,2_ = *α_i_ – α_2_***. We defined the dispersions around the centroids *α*_1_ and *α*_2_ as *β*_1/2_ = MAD((***x***_*i*∈*Y*_1/2__.***γ*_1,2_**)^2^), where. is scalar product of the two vectors, and the normalized distance is:

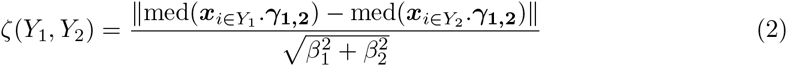

We then iteratively merged all clusters (*Y_i_, Y_j_*) such that *ζ*(*Y_i_,Y_j_*) ≤ *σ*_similar_. At the end of the clustering, every cluster with less than 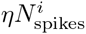 was discarded. In all the manuscript, we used 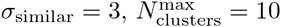, and *η* = 0.005.

#### Template estimation

At the end of the clustering phase, pooling the clusters obtained from every electrode, we obtained for every cluster *Y_k_* a list of spike times 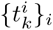. We computed the first component from the raw data as the point-wise median of all the waveforms belonging to the cluster: 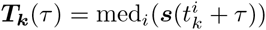. Note that ***T_k_*** is only different from zero on the electrodes close to its peak (see fig. 1D). This information is used internally by the algorithm to save templates as sparse structures. Moreover, during template estimation, we limited the number of spike times per template to a maximal value of 500 to avoid memory saturation. To enhance the compression level of the template ***T_k_***, we set to 0 all the channels *j* where |***T_k_***(*t*)| < *θ_j_*, if *θ_j_* is the detection threshold on channel *j*. This allowed us to remove channels without discriminant information, and to increase the sparsity of the templates. Once the template first component ***T_k_*** had been extracted, we computed its minimal and maximal amplitudes 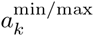 based on data ***x**_i∈Y_k__* used during the clustering. If 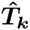 is the normalized template, such that 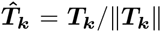, we computed

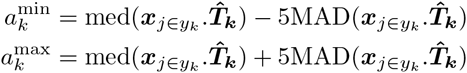

Those boundaries are used during the template matching phase (see below). Finally, we computed the projection of all snippets in the space orthogonal to the first component: 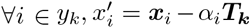, with 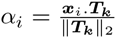. The *x*′ are the projections, and from that we computed the second component of the template, ***U_k_*** (*t*) as the direction of largest variance that is orthogonal to the average waveform ***T_k_**(t)* (i.e. the first principal component).

#### Removing redundant templates

To remove redundant templates that may be present in the dictionary because of the divide and conquer approach (for example a neuron in between two electrodes would give rise to two very similar templates on two electrodes), we computed for all pairs of templates in the dictionary *CC*_max_(*i, j*) = max_*t*_*CC*(***T_i_, T_j_***), where *CC* stands for normalized Cross-Correlation. If *CC*_max_(*i, j*) ≥ *cc*_similar_, we considered those templates to be equivalent and they were merged. In all the following, we used *cc*_similar_ = 0.975. Note that we are computing the Cross-Correlations between normalized templates, such that two templates that have the same shape but different amplitudes are merged. Similarly, we searched if any template ***T_k_*** could be explained as a linear combination of two templates in the dictionary. If we could find ***T_i_*** and ***T_j_*** such that *CC*(***T_k_, T_i_ + T_j_***) ≥ *cc*_similar_, *T_k_* was considered to be a mixture of two cells, and was removed from the dictionary.

### Template matching

At the end of this “template-finding” phase, we have found a dictionary of templates (***T, U***). We now need to reconstruct the signal ***s*** by finding the amplitudes coefficients *a_ij_* and *b_ij_* described in Equation 1. Note that most *a_ij_* and *b_ij_* in this equation are equal to 0. For the other ones, most *a_ij_* values are around 1, because a spike usually appears on electrodes with an amplitude close to the average first component *T*. In this template matching step, all the other parameters have been determined by template extraction and spike detection, so the purpose is only to find the values of these amplitudes. To do so, we used an iterative greedy approach to estimate the *a_ij_* for each subgroup *t_i_*, which bears some similarity to the matching pursuit algorithm [Mallat and Zhang, 1993]. The fitting was performed in blocks of putative spike times, {*t_i_*}, that were successively loaded in memory. Such an approach allowed us to easily split the load linearly among several processors. Each block of raw data ***s*** was loaded and processed according to the following steps during the template-matching phase:

1. Estimate the normalized scalar products ***s***(*t*) · ***T**_j_*(*t* – *t_i_*) for each template *j* and putative spike time *t_i_*, for all the *i* and *j* in the block of raw data.
2. Choose the (*i, j*) pair with the highest scalar product, excluding the pairs (*i, j*) which have already been tried and the *t_i_*’s already explored (see below).
3. Set *a_ij_* equal to the amplitude value that best fits the raw data: 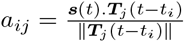.
4. Check if the *a_ij_* amplitude value is between 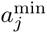 and 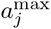.
5. If so, then accept this value, subtract the scaled template from the raw data: s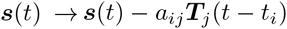. Then set *b_ij_* equal to the amplitude value that best fits the raw data with ***U_j_***, and subtract it too.
6. Return to step 1 to re-estimate the scalar products on the residual.
7. Otherwise, increase by one *n_i_*, which counts the number of times any template has been rejected for spike time *t_i_*. If *n_i_* reaches *n*_failures_ = 3, label this *t_i_* as “explored”. If all t_i_ have been explored, quit the loop. Otherwise return to step 1 and iterate.
8. The parameters of the algorithm were the amplitude thresholds a^min^ and a^max^, computed as described in the Template Estimation section.

### Semi-automated merging

To speed up the manual stage where a human operator has to review all the templates (and thus putative neurons) and decide which one are duplicated or truncated, we developed a dedicated Graphical User Interface (GUI). Such a GUI is especially useful when the number of detected neurons is important, which is the case with the new generation of dense probes. Templates likely to be merged are templates that look alike, and such that the combined cross-correlogram between the two cell’s spike trains show a clear dip near 0ms, indicating that the merged spike trains do not show any refractory period violation (fig 3D, E). To speed up considerably the time spent in processing those datasets, and reducing the inherent variabilities of a human interaction on the sorting process, we provided, for all pairs of templates, a mathematical quantification of the similarity between templates, and of the dip in the cross-correlogram. In all the following *CC_i,j_(t)* represents the Cross Correlogram between the spike times of ***T_i_*** and those of ***T_j_***. For the template similarity, we computed, for every pair of templates (***T_i_, T_j_***), *α*(***T_i_, T_j_***) = max_*t*_*CC_i,j_*(*t*). To quantify the dip in the cross-correlogram, we computed the cross-correlogram *CC_i,j_* (*t*) between the spikes of ***T_i_*** and those of ***T_j_*** and a control version 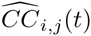 obtained by keeping the spike times from ***T_i_*** but reversing in time those from ***T_j_***. We then computed the average value over time of 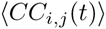 and 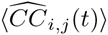 for *t* ≤ *τ*_0_. In the following we used *τ*_0_ = 2ms. This value will depend on the dataset and can be chosen by the user in our Graphical User Interface. From that, we computed the normalized value:

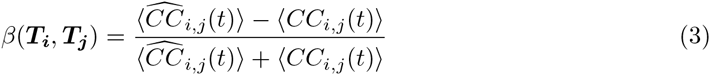

The dip measure *β*(***T_i_, T_j_***) is termed normalized CC metric in fig. 3D,E. It is compared to 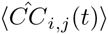, termed Reverse CC in the same figure. For a better visualization, in the insets of fig. 3D,E, the cross-correlograms are such that the mean over time is equal to 1.

For two templates, *α*(***T_i_, T_j_***) allowed us to quantify how similar they are, while *β*(***T_i_, T_j_***) gave us an insight about how strong is the dip in the cross-correlogram between their spike times. To guide the human into the process of merging pairs, we then showed in 2D plots all pairs of templates as function of *α*(***T_i_, T_j_***) and *β*(***T_i_, T_j_***). In such a space, the user can quickly define at once a whole set of pairs that need to be merged. After merging, quantities *α* and *β* are re-computed, and the workflow can keep going until the user decides to stop merging (see fig. 3).

### Simulated ground truth tests

Injection of artificial templates In order to assess the performance of the algorithm, we injected new templates in real datasets (see fig. 4 A-D). To do so, we ran the algorithm on a given dataset, and obtain a list of putative templates 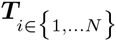 Then, we randomly selected some of those templates ***T_i_*** and shuffled the list of their channels, before injecting them elsewhere in the datasets at controlled firing rates [Harris et al., 2000, Rossant et al., 2016, Kadir et al., 2014]. This allowed us to properly quantify the performance of the algorithm. In order not to bias the clustering, when a template ***T_i_*** was selected and shuffled as a new template 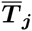 centered on a new channel *j*, we ensured that the injected template was not too similar to one that would already be in the data: 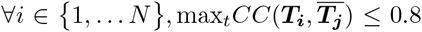. Before being injected, 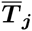 was normalized it such that 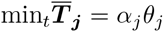. Therefore, the normalized amplitude *α_j_* of 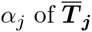, was the relative amplitude, expressed as function of *θ_j_*, the detection threshold on the electrode where the template is peaking. If *θ_j_* ≤ 1, it means that the template has the same height than the noise level, and its spikes should not be detected; if *α* ≥ 1 the spikes should be detected.

#### Injection of correlated templates

In fig. 4E, we selected a particular template ***T***, and kept constant the part on the electrode *i* where it was peaking the most, while shuffling all its neighboring channels. By doing so, we obtained 5 distinct copies of the template 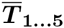, representing 5 neurons whose soma would be located next to the same channel *i*. The normalized amplitudes of those templates were equal to 2. To demonstrate how the code could resolve the problem of overlapping spikes, we injected them into the data such that they were all firing at 10Hz, but with a controlled correlation coefficient *c* that could be varied (using a Multiple Interaction Process [Kuhn et al., 2003]). This parameter *c* allowed us to quantify the percentage of pairwise correlations recovered by the algorithm for overlapping spatio-temporal templates.

#### Generation of artificial spike trains

To illustrate the meta-merging GUI, we generated artificial spike trains where we can have a ground truth for which pairs should be merged. We generated 10 spikes trains *ω_i_* at 10Hz with a correlation coefficient *c* = 0.01 (using a Multiple Interaction Process [Kuhn et al., 2003]), and a refractory period of *τ*_ref_ = 5ms. Then, for every spike trains *ω_i_*, we split it in 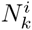 sub spike trains, with 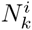 uniformly drawn between [1, 5]. This gave us a number of cells between 10 and 50 (see fig. 3D). The same procedure is repeated in fig. 3E, but starting from 10 neurons obtained after a real sorting session from the rat retina.

#### Estimation of false positive and false negative

To quantify the performance of the algorithm, we matched the spikes recovered by the algorithm to the real ground-truth spikes (either synthetic or obtained with juxta-cellular recordings). A spike was considered to be a match if it had a corresponding spike in the ground truth at less than 2ms. Spikes in the ground-truth datasets that had no matches in the spike sorting results in a 2 ms window were labeled as “False Negative”, while those that are not present while the algorithm detected a spike were “False Positive”. False negative rate was defined as the number of false negative divided by the number of spikes in the ground truth recording. False positive rate was defined as the number of false positive divided by the number of spikes in the spike train extracted by the algorithm. In the paper, the error is defined as mean of the False negative and the False positive rates (see fig. 4, 5). Note that to take into account the fact that a ground-truth neuron could be split in several templates at the end of the algorithm, we always compared the ground-truth cells with the combination of templates that minimized the error.

#### Theoretical estimate

To quantify the performance of the software with real ground-truth recordings (see fig. 5), we computed the Best Ellipsoidal Error Rate (BEER), as described in [Harris et al., 2000]. This BEER estimate gave an upper bound on the performance of any clustering-based spike sorting method using elliptical cluster boundaries. After thresholding and feature extraction, snippets were labeled according to whether or not they contained a true spike. Half of this labeled data set was then used to train a perceptron whose decision rule is a linear combination of all pairwise products of the features of each snippet. If *x_i_* is the *i*-th snippet, projected in the feature space, then the optimized function *f*(*x*) is:

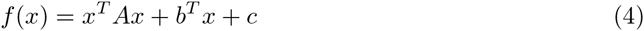

We trained this function *f* by varying *A, b* and *c* with the objective that *f(x)* should be 1 for the ground truth spikes, and −1 otherwise. These parameters were optimized by a stochastic gradient descent with a regularization constraint. The resulting classifier was then used to predict the occurrence of spikes in the snippets in the remaining half of the labeled data. Only the snippets where *f(x)* > 0 were predicted as true spikes. This prediction provided an estimate of the false negative and false positive rates for the BEER estimate. The mean between the two was considered to be the BEER error rate.

#### Decimation of the electrodes

In order to increase the number of data points for the comparison between our sorting algorithm and the non-linear classifiers defined by the BEER metric (see fig. 5), we ran the analysis several times on the same neurons, but removing some electrodes, to create recordings at a lower electrode density. We divided by a factor 2 or 4 the number of electrodes in the 252 in *vitro* Multi-Electrode Array or the 128 in vivo silicon probe.

### Implementation and Source Code

The code is a pure python package, based on the python wrapper for the Message Passing Interface (MPI) library [Dalcin et al., 2011] to allow parallelization over distributed computers, and is available with its full documentation at http://spyking-circus.rtfd.org. Results can easily be exported to the kwik or phy format [Rossant and Harris, 2013]. All the datasets used in this manuscript will also be available on-line, for testing and comparison with other algorithms.

## Acknowledgments

We would like to thank Steve Baccus and Sami El Boustani for insightful discussions. We also would like to thanks Kenneth Harris, Cyrille Rossant and Nick Steimetz for feedbacks and the help with the interface to the phy software.

